# ATF4 and mTOR regulate metabolic reprogramming in TGF-β-treated lung fibroblasts

**DOI:** 10.1101/2024.06.12.598694

**Authors:** Kun Woo D Shin, M Volkan Atalay, Rengul Cetin-Atalay, Erin M O’Leary, Mariel E Glass, Jennifer C Houpy Szafran, Parker S Woods, Angelo Y Meliton, Obada R Shamaa, Yufeng Tian, Gökhan M. Mutlu, Robert B. Hamanaka

**Affiliations:** Department of Medicine, Section of Pulmonary and Critical Care Medicine, The University of Chicago, Chicago, IL 60637; Middle East Technical University, Ankara, Turkey

## Abstract

Idiopathic pulmonary fibrosis is a fatal disease characterized by the TGF-β-dependent activation of lung fibroblasts, leading to excessive deposition of collagen proteins and progressive replacement of healthy lung with scar tissue. We and others have shown that fibroblast activation is supported by metabolic reprogramming, including the upregulation of the *de novo* synthesis of glycine, the most abundant amino acid found in collagen protein. How fibroblast metabolic reprogramming is regulated downstream of TGF-β is incompletely understood. We and others have shown that TGF-β-mediated activation of the Mechanistic Target of Rapamycin Complex 1 (mTORC1) and downstream upregulation of Activating Transcription Factor 4 (ATF4) promote increased expression of the enzymes required for *de novo* glycine synthesis; however, whether mTOR and ATF4 regulate other metabolic pathways in lung fibroblasts has not been explored. Here, we used RNA sequencing to determine how both ATF4 and mTOR regulate gene expression in human lung fibroblasts following TGF-β. We found that ATF4 primarily regulates enzymes and transporters involved in amino acid homeostasis as well as aminoacyl-tRNA synthetases. mTOR inhibition resulted not only in the loss of ATF4 target gene expression, but also in the reduced expression of glycolytic enzymes and mitochondrial electron transport chain subunits. Analysis of TGF-β-induced changes in cellular metabolite levels confirmed that ATF4 regulates amino acid homeostasis in lung fibroblasts while mTOR also regulates glycolytic and TCA cycle metabolites. We further analyzed publicly available single cell RNAseq data sets and found increased expression of ATF4 and mTOR metabolic targets in pathologic fibroblast populations from the lungs of IPF patients. Our results provide insight into the mechanisms of metabolic reprogramming in lung fibroblasts and highlight novel ATF4 and mTOR-dependent pathways that may be targeted to inhibit fibrotic processes.

## INTRODUCTION

Idiopathic Pulmonary Fibrosis (IPF) is a progressive, fatal disease, which has a median survival of 3.5 years and affects approximately 150,000 people in the United States (1, 2). A defining feature of IPF is the Transforming Growth Factor-β (TGF-β)-dependent activation of lung fibroblasts, leading to the excessive secretion of extracellular matrix proteins, including collagen (3–6). Activated fibroblasts are the primary cells responsible for the structural remodeling and impairment of lung function characteristic of IPF and thus represent a key therapeutic target for the treatment of the disease (6–8).

Metabolic reprogramming has emerged as a key regulator of fibroblast activation and is increasingly studied as a target of therapeutic intervention for IPF (9–12). Treatment of lung fibroblasts with the profibrotic cytokine TGF-β increases both glycolytic rate and mitochondrial respiration, the inhibition of which prevents matrix production (13–17). We and others have found that TGF-β increases *de novo* glycine synthesis in lung fibroblasts (11, 15, 16, 18–20). Glycine is a non-essential amino acid that constitutes over one third of the primary structure of collagen protein and we have shown that inhibition of this pathway reduces TGF-β-induced collagen protein production *in vitro*, and ameliorates bleomycin-induced lung fibrosis *in vivo* (15, 16, 18).

Upregulation of *de novo* glycine synthesis downstream of TGF-β requires ATF4 (Activating Transcription Factor 4)-dependent transcriptional activation of the metabolic enzymes of the serine, glycine, one-carbon (SGOC) pathway, which converts the glycolytic intermediate 3-phosphoglycerate to the amino acids serine and glycine (17, 19). Activation of ATF4 is regulated by the Mechanistic Target of Rapamycin Complex 1 (mTORC1) which promotes ATF4 protein translation (21–23). mTOR has also been shown to regulate metabolic transcriptional programs induced by other transcription factors including HIF-1α (Hypoxia Inducible Factor-1 α), SREBP1/2 (Sterol Regulatory Element Binding Protein 1/2), PGC-1α (Peroxisome Proliferator-Activated Receptor-Gamma Coactivator 1α), and TFEB (Transcription Factor EB) (24–32). How ATF4 and mTOR regulate metabolism in lung fibroblasts is incompletely understood.

Here, we used RNAseq to determine the contribution of ATF4 and mTOR to gene expression and metabolic reprogramming downstream of TGF-β. We found that in addition to glycine biosynthetic enzymes, ATF4 regulates expression of amino acid transporters and aminoacyl-tRNA synthetases. Using gas chromatography-mass spectrometry, we show that ATF4 is required for increased cellular levels of glycine, cysteine, asparagine, and branched chain amino acids downstream of TGF-β. mTOR is required for expression of ATF4 target genes and additionally promotes expression of glycolytic enzymes and subunits of the mitochondrial electron transport chain. mTOR inhibition prevented TGF-β-mediated increases in cellular amino acid levels, but also increases in TCA cycle metabolites and lactate. Finally, we analyzed the expression of ATF4 and mTOR target genes in publicly available single cell RNAseq data sets, providing evidence for metabolic reprogramming in fibroblasts in pulmonary fibrosis *in vivo*.

## MATERIALS AND METHODS

### Fibroblast Culture

Normal human lung fibroblasts (HLFs) (Lonza) were cultured as previously described (15). Cells were serum starved in DMEM (Gibco) containing 0.1% bovine serum albumin (BSA), 5.5mM glucose, 2 mM glutamine, and 1mM pyruvate for 24 hours prior to treatment with TGF-β (1ng/mL, Peprotech). TORIN1 (R&D Systems), was added 30 minutes prior to TGF-β.

### siRNA Knockdowns

For siRNA knockdowns, 1×10^6^ normal NHLFs were transfected with 250 pmol ON-TARGETplus siRNA (Dharmacon). Cells were plated on 10cm dishes for 24 hours and then replated for experiments as above. Dharmacon product numbers are: nontargeting siRNA-D-001810-01, siATF4 1-J-005125-10, siATF4 2-J-005125-12.

### RNA Isolation and Quantitative PCR

RNA as isolated using the GenElute Total RNA Purification Kit (Sigma) and reverse transcribed using iScript Reverse Transcription Supermix (Bio-Rad). Quantitative mRNA expression was determined by real-time RT-PCR using ITaq Universal SYBR Green Supermix (Bio-Rad). For a list of primers used, see online supplementary material.

### RNAseq Analysis

Total RNA isolated as above was sequenced on an Illumina NovaSEQ6000 at the University of Chicago Genomics Core Facility (100bp single end). Sequence qualities of generated FASTQ files were assessed using FastQC. Transcript expression was quantified using Salmon v1.10.1 (33). The Salmon index was created with the GENCODE (Human Release 45, GRCh38.14) quantification was performed in quant mode using default parameters. Gene abundances were computed using the R package txiimport v.1.28.0 (34). Non-coding RNAs were filtered out from quantified data. Batch effects were corrected using the sva package v.3.48.0 (35). Differential gene expression analysis was performed using the edgeR package v. 3.42.4 quasi-likelihood F model (36). Differential gene expression was considered significant for genes with an FDR-adjusted p-value ≤0.05. Gene set enrichment alalysis was performed on Hallmark pathways from MsigDB using the clusterProfiler package v.4.10.1 and GSEA software and on Reactome pathways using the Reactome PA package v.1.46.0 (37–39). Transcript Factor Enrichment Analysis was performed using the DoRothEA package v.1.12.0 (40). All packages were run in RStudio (2023.06.2+561) with R version 4.3.1.

### Gas Chromatography Mass Spectrometry

Normal HLFs grown for 48 hours in the presence or absence of TGF-β were washed in blood bank saline and metabolites were extracted in 600μL ice cold 80% MeOH. Samples were derivatized as we have previously (41) and analyzed with an 8890 gas chromatograph with an HP-5MS column (Agilent) coupled with a 5977B Mass Selective Detector mass spectrometer (Agilent) as we have previously (41). Peak ion chromatograms for metabolites of interest were extracted at their specific *m/z* with Mass Hunter Quantitative Analysis software (Agilent). Ions used for quantification of metabolite levels were as follows: Glycine *m/z* 246, Asparagine *m/z* 417, Leucine *m/z* 302, Isoleucine *m/z* 302, Valine *m/z* 288, Glutamate *m/z* 432, Proline *m/z* 258, Methionine *m/z* 320, Lactate *m/z* 261, α-ketoglutarate *m/z* 346, Succinate *m/z* 289, Malate *m/z* 419.

### Analysis of scRNAseq Data sets

Preanalyzed data from Habermann *et al* (42) were downloaded from GitHub (https://github.com/tgen/banovichlab/tree/master/pulmonary_fibrosis/10x_scRNA-Seq_2019). This dataset is also publicly available at GEO: GSE135893. scRNA-seq data were processed and analyzed using Seurat v5 package in R version 4.4.0. We excluded cells with fewer than 250 detected genes or larger than 20% mitochondrial genes. Seurat v5 was used to perform dimensionality reduction, clustering, and visualization. Recursive clustering analysis of subpopulations was conducted to improve the granularity of cell annotations and to obtain fibroblast cells. Cell-type annotation of fibroblasts was performed based on markers defined by Habermann. Visualization of the cells and clusters on a 2D map was performed with uniform manifold approximation and projection (UMAP). Dot plots and UMAP plots overlaid with gene expression levels were generated using Seurat.

### Statistical analysis

qRT-PCR and metabolomic data were analyzed in Prism 10 (GraphPad Software, Inc). All data are shown as mean ± standard error of the mean (SEM). Significance was determined by one-way or two-way ANOVA using Tukey’s correction for multiple comparisons. * P < 0.05, ** P < 0.01, *** P < 0.001.

## RESULTS

### TGF-β regulates the expression of metabolic enzymes in lung fibroblasts

To determine how ATF4 and mTOR regulate cellular metabolism in human lung fibroblasts (HLFs) we 1) knocked down *ATF4* using siRNA, or 2) treated cells with the mTOR kinase inhibitor TORIN1 as we have previously described (17). Cells were treated with TGF-β for 24 hours or left untreated, after which cells were harvested and RNA extracted for sequencing. Principal component analysis shows that gene expression of untreated cells cluster together and that the largest effect on mRNA expression was due to TGF-β exposure **(Fig. 1A)**. As expected, the effect of mTOR inhibition was relatively greater than that of *ATF4* knockdown. We found that TGF-β induced the expression of 634 genes in HLFs, while the expression of 568 genes was repressed (Log2FC ≥ 1, FDR ≤ 0.05) **(Fig. 1B)**. Hallmark Pathway Analysis on genes differentially expressed between untreated and TGF-β treated cells showed that in addition to hallmarks of TGF-β signaling, myogenesis, and epithelial mesenchymal transition, genes associated with hypoxia responses, glycolysis, mTORC1 signaling, and the unfolded protein response (UPR) were also highly induced by TGF-β **(Fig. 1C)**. Among the genes associated with mTORC1 signaling and the UPR were the amino acid biosynthetic enzymes involved in synthesis of glycine (*PHGDH*, *PSAT1*, *PSPH*, *SHMT2*, and *MTHFD2*), proline (*ALDH18A1*), and asparagine (*ASNS*). Amino acid transporters (*SLC1A4*, *SLC7A5*) were also enriched in these pathways **(Fig. 1D)**. Genes associated with glycolysis and responses to hypoxia included the glucose transporter *SLC2A1*, glycolytic enzymes (*PFKL* and *PFKP*), lactate dehydrogenase (*LDHA*), and lactate transporter (*SLC16A3*) (Figure 1D). We also found significant downregulation of genes involved in fatty acid metabolism (*CPT1A*, *FASN*). We found that *PSAT1*, *MTHFD2*, *SLC7A5*, and *ASNS* were among the top 50 genes regulated by TGF-β **(Fig. 1E)**. These findings suggest that metabolic reprogramming is one of the key functions of TGF-β signaling in lung fibroblasts.

**Figure 1.**
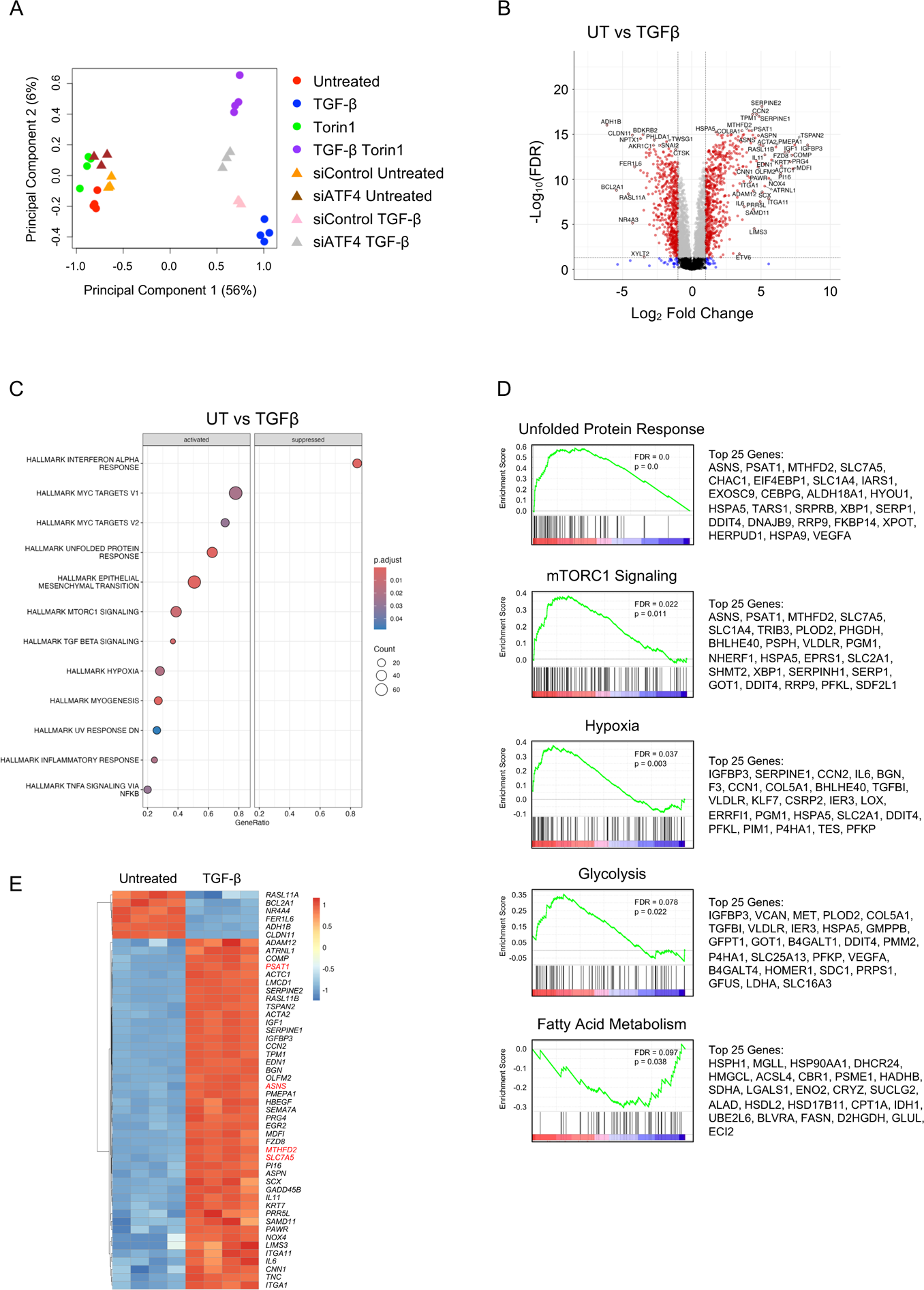
TGF-β promotes metabolic reprogramming in human lung fibroblasts. **(A)** Multidimensional scaling plot of differentially expressed genes (DEGs) in HLFs cultured in the presence or absence of TGF-β (1ng/mL). Cells were either transfected with a nontargeting siRNA or siRNA targeting ATF4. Alternatively, cells were cultured in the presence or absence of the mTOR kinase inhibitor TORIN1 (125nM) **(B)** Volcano plot of -Log10(FDR) vs. Log2 fold change for DEGs between untreated and TGF-β-treated HLFs **(C)** Significantly activated and suppressed Molecular Signatures Database (MSigDB) Hallmark pathways enriched in DEGs between untreated and TGF-β-treated HLFs. **(D)** Enrichment plots of significantly enriched metabolism-related Hallmark pathways regulated by TGF-β treatment in HLFs. The top 25 enriched genes in each pathway are indicated. **(E)** Heatmap analysis of the top 50 significant DEGs between untreated and TGF-β-treated HLFs.

### ATF4 regulates genes involved in amino acid biosynthesis and tRNA aminoacylation in TGF-β-treated lung fibroblasts

To determine how ATF4 regulates the response to TGF-β we analyzed DEGs between TGF-β-treated fibroblasts that had been transfected with either non-targeting siRNA or siRNA targeting *ATF4*. We found a total of 14 genes that are significantly regulated by ATF4 in lung fibroblasts (Log2FC ≥ 1, FDR ≤ 0.05) **(Fig. 2A)**. All 14 DEGs were downregulated in ATF4 knockdown cells compared with control siRNA-transfected cells **(Fig. 2B)**. Gene set enrichment analysis (GSEA) using Reactome on all genes regulated by ATF4 with FDR < 0.05 **(Fig. S1A)**, showed that these DEGs largely segregate into two groups-one that is highly enriched for genes involved in amino acid biosynthetic enzymes, and another enriched for genes involved in stress responses **(Fig. 2C, 2D)**. As we have previously shown, genes required for *de novo* synthesis of glycine and one carbon metabolism were represented among the ATF4 target genes (*PHGDH*, *PSAT1*, *MTHFD2*, *ALDH1L2*) (15, 18, 41). Also present were *ASNS* and amino acid transporters (*SLC1A4*, *SLC7A5*, *SLC6A9*) **(Fig. 2A)**. Analysis of genes that were significantly regulated by ATF4, but with a lower log2FC revealed that ATF4 also regulates the expression of aminoacyl tRNA synthetases (*AARS1*, *IARS1*, *GARS1*, *WARS1*) in HLFs, suggesting a major role of ATF4 in promoting amino acid homeostasis and protein synthesis during fibrotic responses **(Fig. S1A)**. Genes associated with stress responses such as *HERPUD1*, *DDIT4*, and *CHAC* were also sensitive to ATF4 knockdown, likely due to low levels of ER stress that are induced by TGF-β as we have previously shown (17). We confirmed the effect of *ATF4* knockdown on these target genes using two independent siRNAs **(Fig. S1B)**.

**Figure 2.**
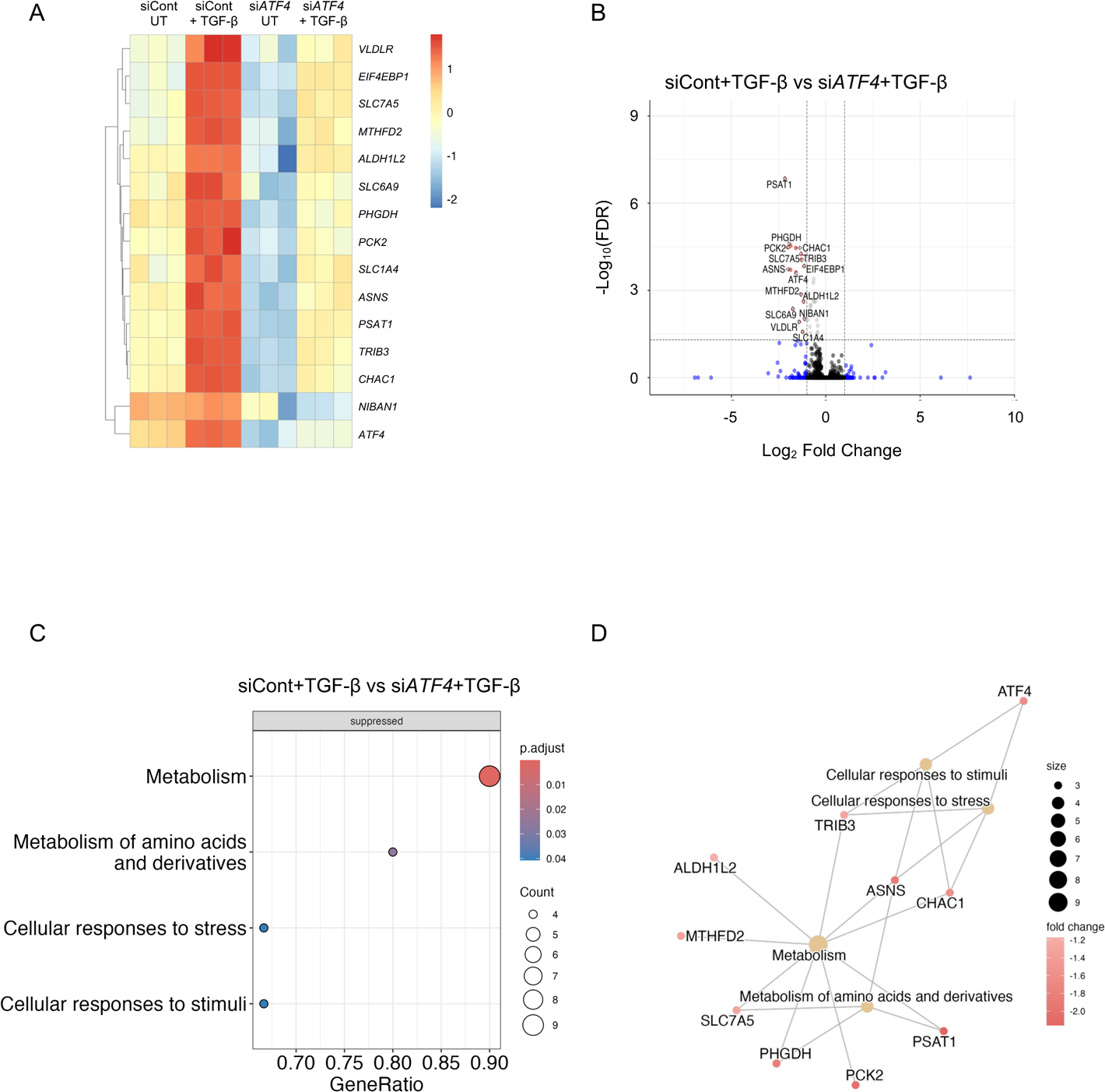
ATF4 regulates the expression of amino acid transporters and biosynthetic enzymes in HLFs. **(A)** Heatmap analysis of significant DEGs (log2FC≥1) between TGF-β-treated HLFs with either control or ATF4 knockdown. **(B)** Volcano plot of -Log10(FDR) vs. Log2 fold change for DEGs between control and ATF4 knockdown HLFs treated with TGF-β. **(C)** Significantly suppressed Reactome pathways enriched in DEGs between control and ATF4 knockdown HLFs treated with TGF-β. **(D)** CNET plot of Reactome pathways enriched in DEGs between control and ATF4 knockdown HLFs treated with TGF-β

### mTOR regulates multiple metabolic pathways in TGF-β-treated lung fibroblasts

To determine how mTOR regulates the response to TGF-β we analyzed DEGs between TGF-β-treated fibroblasts that had been treated with either TORIN1 or vehicle. We found that inhibition of mTOR had a greater effect on TGF-β-induced gene expression than *ATF4* knockdown, with 476 genes differentially regulated between control and TORIN1-treated cells (Log2FC ≥ 1, FDR ≤ 0.05) **(Fig. 3A)**. Unlike ATF4 knockdown, mTOR inhibition led to both increases and decreases in gene expression in TGF-β-treated HLFs (301 genes upregulated, 175 genes downregulated) **(Fig. 3B)**. Hallmark pathway analysis showed that in addition to negative regulation of UPR and mTORC1 signaling hallmarks, genes involved in oxidative phosphorylation and glycolysis were also inhibited by TORIN1 treatment **(Fig. 3C, S2A)**. Hallmark glycolysis genes downregulated by TORIN1 included *LDHA* and the lactate transporter *SLC16A3* **(Fig. 3D)**. Hallmark oxidative phosphorylation genes included subunits of each respiratory complex (Complex I-*NDUFA6*, *NDUFA8*, *NDUFB3*, *NDUFAB6*, *NDUFA1*, *NDUFS8*, *NDUFAB1*, *NDUFA9*, *NDUFB7*). Complex II-*SDHC*. Complex III-*UQCR11*, *UQCR10*. Complex IV-*COX8A*, *COX7A2*, *COX7B*. Complex V-*ATP5F1E*, *ATP5MC3*, *ATP6MF*) **(Fig. 3D)**. These findings are consistent with our previous finding that mTOR inhibition prevents TGF-β-mediated increases in glycolytic rate and mitochondrial oxygen consumption in HLFs (17). We confirmed the effect of TORIN1 on TGF-β-induced expression of glycolytic and respiratory complex genes by qPCR **(Fig. S2B).**

**Figure 3.**
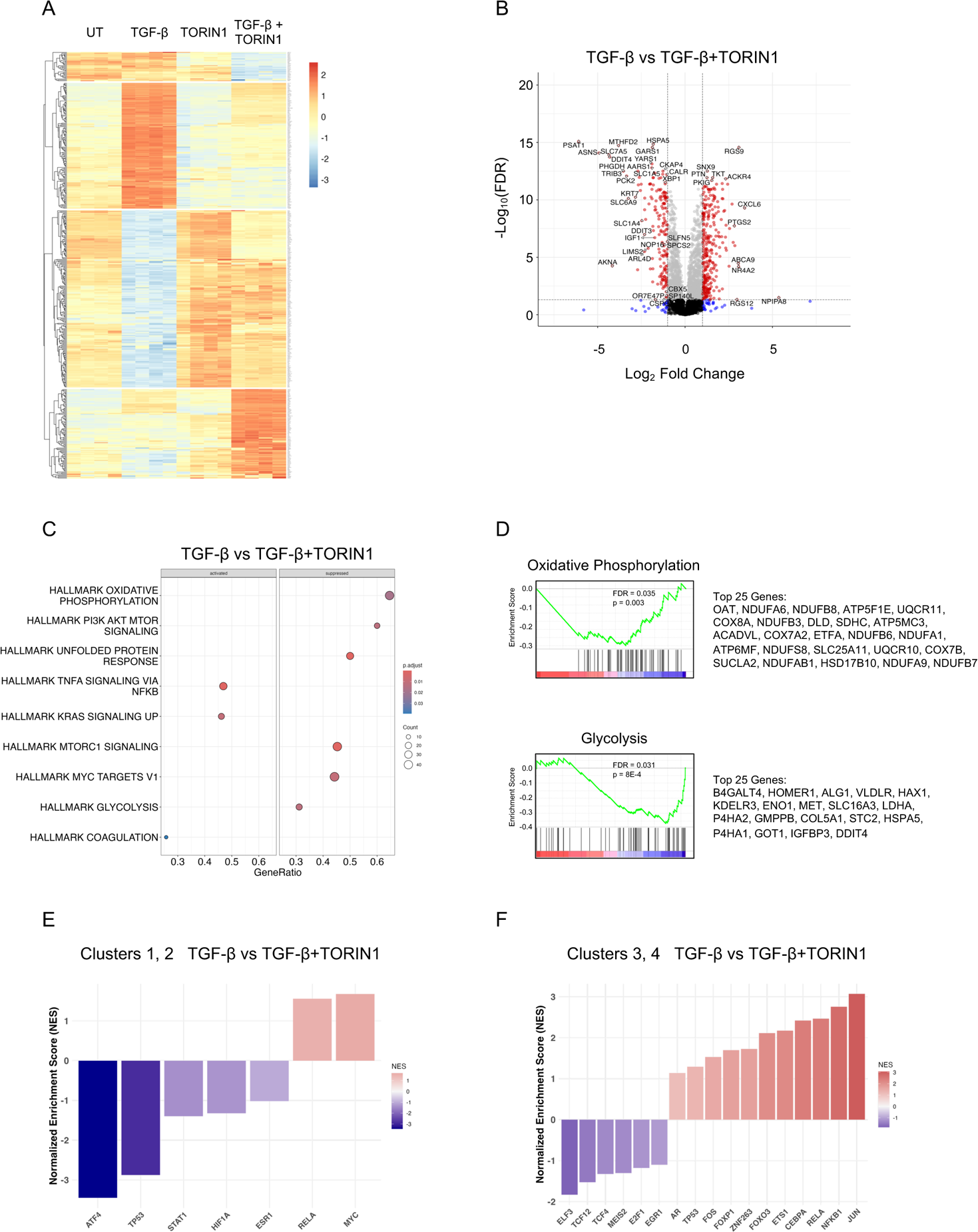
mTOR regulates the expression of genes encoding glycolytic enzymes and subunits of the mitochondrial respiratory chain. **(A)** Heatmap analysis of significant DEGs (log2FC≥1) between TGF-β-treated HLFs cultured in the presence or absence of TORIN1. **(B)** Volcano plot of -Log10(FDR) vs. Log2 fold change for DEGs between TGF-β-treated HLFs cultured in the presence or absence of TORIN1. **(C)** Significantly activated and suppressed MSigDB Hallmark pathways enriched in DEGs between TGF-β-treated HLFs cultured in the presence or absence of TORIN1. **(D)** Enrichment plots of significantly enriched metabolism-related Hallmark pathways regulated by mTOR inhibition in TGF-β-treated HLFs. The top 25 enriched genes in each pathway are indicated. **(E-F)** Transcription factor enrichment analysis using DoRothEA human regulon for TORIN1 (E) downregulated and (F) upregulated genes.

Heatmap analysis of mTOR-sensitive DEGs showed that these genes clustered into 4 groups based on response to TGF-β and to mTOR inhibition **(Fig. 3A)**. Clusters 1 and 2 contained genes that are downregulated by mTOR inhibition with cluster 2 being more sensitive to TGF-β. To determine how transcription factors (TFs) other than ATF4 may be regulated by mTOR inhibition, TF activities were analyzed using DoRothEA TF-target interactions regulons tool. We found that ATF4 and HIF-1α associated genes were among the highly downregulated by mTOR inhibition in Clusters 1 and 2 **(Fig. 3E)**. STAT1 and p53 targets were also highly downregulated by TORIN1. RELA and MYC targets were predicted to be negative regulators of genes inhibited by TORIN1 in Clusters 1 and 2. Analysis of genes that were upregulated by TORIN1 in Clusters 3 and 4, we found that JUN, NFKB1, CEBPA, and FOXO3 were among the TFs most highly repressed by mTOR activity **(Fig. 3F)**. In cancer cells, mTOR has been shown to regulate SREBP-dependent transcription, promoting expression of genes involved in fatty acid synthesis and the pentose phosphate pathway (29). While we did not find SREBP targets to be enriched in our analyses, we queried genes involved in these metabolic pathways to determine how they are regulated in HLFs by TGF-β and mTOR **(Fig. S3)**. We found no consistent regulation of either pathway by mTOR inhibition, suggesting that regulation of SREBP-mediated transcription is not a major factor in TGF-β-mediated activation of lung fibroblasts.

### ATF4 and mTOR regulate cellular amino acid and central carbon metabolites

To determine how the transcriptional programs regulated by ATF4 and mTOR affect cellular metabolite levels, we extracted metabolites from HLFs in the presence or absence of TGF-β with either *ATF4* knockdown or TORIN1 treatment and analyzed by gas chromatography-mass spectrometry. Consistent with a role for ATF4 in regulating glycine and asparagine biosynthesis, we found that TGF-β-mediated increases in these amino acids were attenuated in *ATF4* knockdown cells **(Fig. 4A)**. We also found that ATF4 knockdown cells contained reduced levels of several amino acids that are known to be imported through SLC7A5 (43, 44). These included the branched chain amino acids leucine, isoleucine, and valine as well as tyrosine, phenylalanine, and tryptophan **(Fig. 4A, S4A)**. Other amino acids including glutamate, proline, methionine, histidine, and threonine were not affected by *ATF4* knockdown **(Fig. 4A, S4A)**. Consistent with our previous finding that *ATF4* knockdown does not affect glycolytic flux or mitochondrial oxygen consumption in HLFs, lactate and the TCA cycle intermediates α-ketoglutarate, succinate, malate, and fumarate were not affected by *ATF4* knockdown **(Fig. 4B)**.

**Figure 4.**
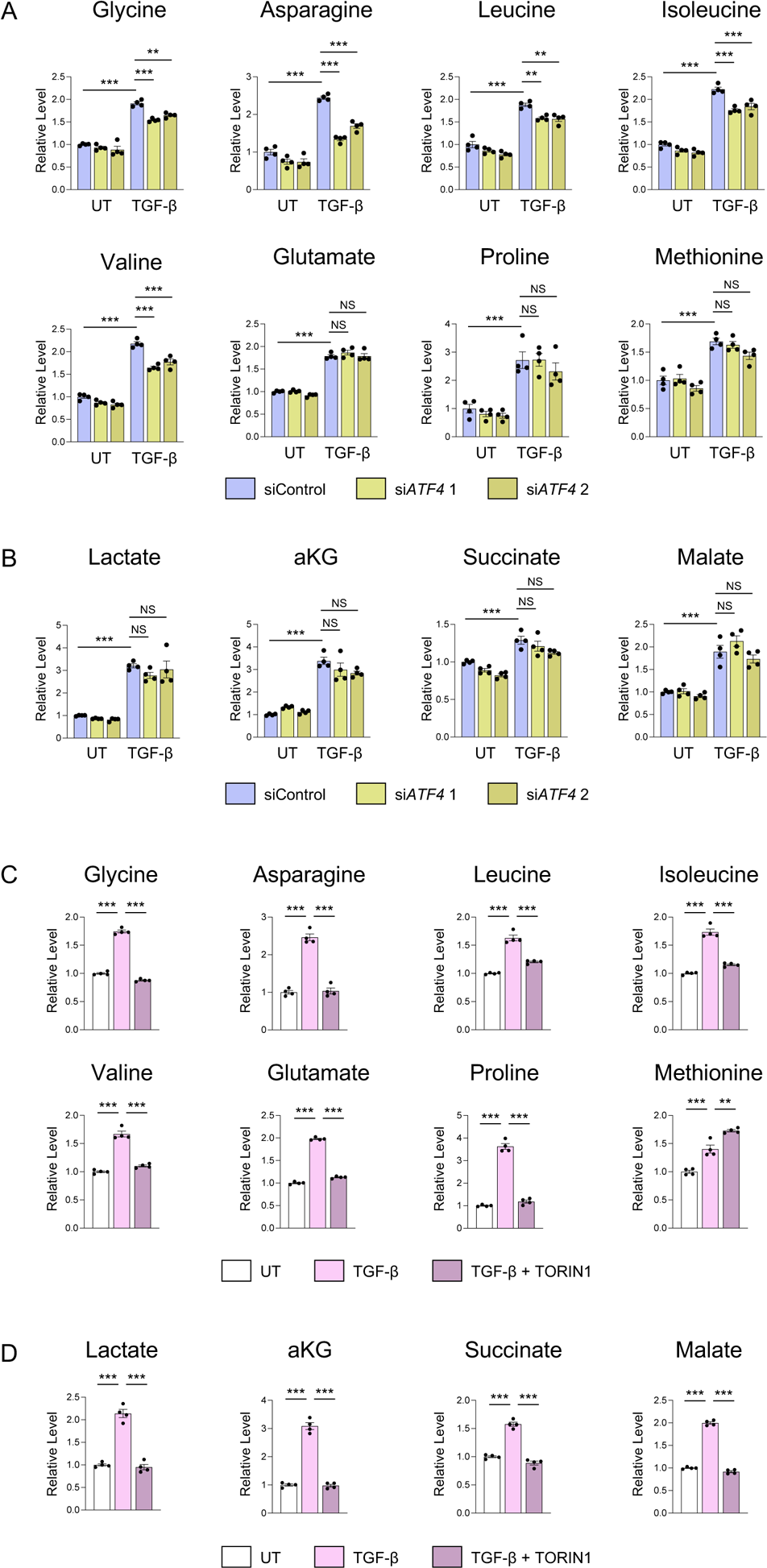
ATF4 and mTOR regulate TGF-β-induced changes in cellular metabolite levels in HLFs. **(A)** Relative cellular levels of the indicated amino acids in HLFs treated with TGF-β for 48 hours. Cells were transfected with either nontargeting siRNA or siRNA targeting ATF4 **(B)** Relative cellular levels of the lactate or the indicated TCA cycle intermediates in HLFs treated with TGF-β for 48 hours. Cells were transfected with either nontargeting siRNA or siRNA targeting ATF4 **(C)** Relative cellular levels of the indicated amino acids in HLFs treated with TGF-β for 48 hours. Cells were cotreated with TORIN1 as indicated. **(D)** Relative cellular levels of the lactate or the indicated TCA cycle intermediates in HLFs treated with TGF-β for 48 hours. Cells were cotreated with TORIN1 as indicated. \**P*<0.05, \*\**P*<0.01, \*\*\**P*<0.001.

mTOR inhibition reduced cellular levels of glycine, asparagine, and SLC7A5-linked amino acids **(Fig. 4C, S4B)**. Cellular glutamate and proline levels were also reduced by TORIN1 treatment while histidine and threonine remained unaffected **(Fig. 4C, S4B)**. In contrast to ATF4 knockdown, our previous findings show that mTOR inhibition prevents TGF-β-mediated inductions in glycolysis and oxygen consumption. Consistent with these findings, we found that TORIN1 reduced TGF-β-mediated increases in lactate, alpha-ketoglutarate, succinate, and malate **(Fig. 4D, S4B)**.

### Expression of ATF4 and mTOR target genes in pulmonary fibrosis patient fibroblasts

Recent studies using single cell RNAseq (scRNAseq) have shed light on the heterogeneity of fibroblast populations in healthy and IPF lungs as well as the changes that occur during disease pathogenesis. To determine whether ATF4 and mTOR-dependent genes are expressed in pathologic cell populations in IPF lungs, we analyzed expression of select genes in a previously published data set from 10 control subjects and 20 patients with pulmonary fibrosis (42). Mesenchymal cells, identified by expression of *COL1A1*, *PDGFRA*, *PDGFRB*, and *LUM* were further subclustered and the fibroblast population (expressing *LUM*, *FBLN2*, and *PDGFRA*) was segregated from smooth muscle, pericytes, and mesothelial cells **(Fig. S5A-S5B)**. After reclustering of the fibroblast population, we identified a central fibroblast population that contained the most cells from control samples **(Fig. S5C, S5D)**. Disease-associated populations were annotated as defined by Habermann *et al* **(Fig. S5E)**. This included myofibroblast and PLIN2^+^ populations that were expanded in disease lungs as well as a disease-specific HAS1^High^ population **(Fig. 5A)**.

**Figure 5.**
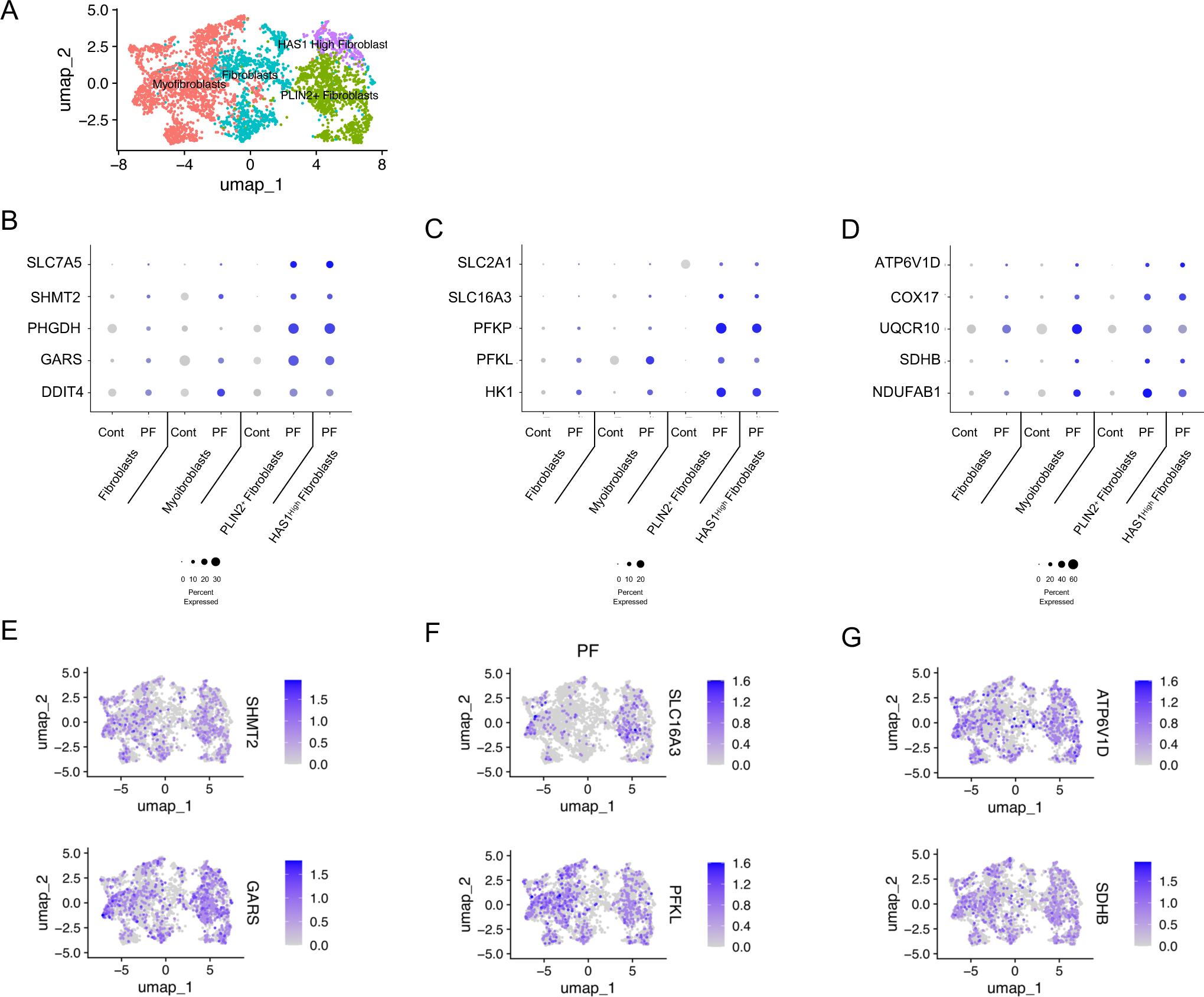
ATF4 and mTOR target genes are expressed in pathologic fibroblast populations of pulmonary fibrosis patients. **(A)** Clustering of fibroblasts from 10 control lungs and 20 lungs from pulmonary fibrosis patients. Populations were assigned as defined by Habermann et all (42). **(B-C)** Dot plot representation of the expression of select (B) ATF4 target genes, (C) glycolysis-related genes, and (D) respiratory chain subunits in fibroblast subpopulations and stratified by disease state. **(E-G)** UMAP projection of expression of select (E) ATF4 target genes, (F) glycolysis-related genes, and (G) respiratory chain subunits in fibroblasts from pulmonary fibrosis lungs.

We examined the expression of ATF4 targets **(Fig. 5B)** as well as glycolysis **(Fig. 5C)** and oxidative phosphorylation **(Fig. 5D)** associated genes. We found that within each population, expression of these genes was elevated in cells from disease lungs compared with non-disease controls. Furthermore, gene expression was generally highest in the PLIN2^+^ and HAS1^High^ populations and lowest in other fibroblast populations. Examining gene expression in disease cells by UMAP plot showed heterogeneity of gene expression across populations. Within fibroblast populations, gene expression was generally highest in cells co-expressing PI16, a marker for adventitial fibroblasts **(Fig. S5F)**. Gene expression in myofibroblast populations showed highest levels in CTHRC1^High^ cells, which have the highest *COL1A1* expression **(Fig. S5F)**. Together these findings suggest that similar metabolic reprogramming occurs during fibrotic responses in fibroblasts both *in vitro* and *in vivo*.

## DISCUSSION

Metabolic reprogramming in lung fibroblasts is an emerging mechanism of the pathogenesis of pulmonary fibrosis (9–12). We and others have previously demonstrated that activation of mTOR and downstream activation of ATF4 are required for induction of enzymes required for *de novo* glycine synthesis, which supports the production of collagen proteins (17, 19). How these regulators affect expression of other genes as well as metabolite levels in lung fibroblasts is poorly understood. Using RNA sequencing, we queried mTOR and ATF4-dependent gene expression in TGF-β-treated lung fibroblasts. We found that both mTOR and ATF4 contribute to amino acid homeostasis in HLFs; however, mTOR is also a key regulator of glycolytic and mitochondrial metabolism.

ATF4 is best known for its activation by the integrated stress response (ISR) which leads to phosphorylation of eukaryotic initiation factor 2α and selective translation of *ATF4* mRNA (45). Our previous findings have shown that TGF-β treatment results in activation of the ER stress response in HLFs, leading to increased expression of *CHOP* mRNA and increased expression and splicing of *XBP1* mRNA (17). Consistent with activation of the ISR, we found ATF4-dependent expression of stress-related genes such as *HERPUD1*, *DDIT4*, and *CHAC* in TGF-β-treated HLFs. How the other branches of the ER stress response regulate TGF-β-treated gene expression in HLFs is yet to be determined; however, it is likely that activation of this pathway promotes expansion of the ER to support matrix protein production.

In addition to regulation by stress, ATF4 is induced downstream of growth factor signaling, through activation of mTORC1, allowing cells to coordinate amino acid biosynthesis and uptake capacity with growth signals (19, 21–23, 46). The exact mechanisms by which mTOR promotes ATF4 activation are incompletely understood; however, eIF2α phosphorylation and thus activation of the ISR is dispensable for this activation. As we have previously shown, enzymes required for production of glycine were highly enriched among the ATF4 target genes (17). In addition to serine-glycine-one carbon metabolic enzymes, we found that ATF4 also regulates the expression of *ASNS*, amino acid transporters, and aminoacyl tRNA synthases in HLFs. Consistent with this, we found that cellular levels of glycine, asparagine, and neutral amino acids linked to SLC7A5 were reduced in *ATF4* knockdown HLFs. These findings suggest that the major role for ATF4 in HLFs is in promoting amino acid homeostasis and tRNA charging to support increased protein production downstream of TGF-β.

mTOR is a master regulator of cellular growth and proliferation, coordinating signals from growth factors and nutrient levels with outputs including protein translation, autophagy, and metabolism. mTOR has been shown to regulate multiple transcriptional programs that regulate metabolic reprogramming in addition to ATF4, including HIF-1α, SREBPs, and PGC-1α, promoting glycolysis, fatty acid metabolism, and mitochondrial biogenesis (24, 25, 28, 29, 47). Our findings show that mTOR is an important transcriptional regulator of glycolytic enzymes and mitochondrial respiratory subunits in HLFs. Consistent with this, we found that mTOR inhibition resulted in reduced cellular levels of lactate as well as TCA cycle intermediates while loss of ATF4 did not. We also observed reduced cellular glutamate and proline levels in TORIN1-treated HLFs. We and others have shown that glutamine metabolism and production of proline are important regulators of collagen production by HLFs (16, 20, 48, 49). How mTOR regulates this process will require further exploration.

Like ATF4, HIF-1α has been shown to be induced downstream of TGF-β and promotes glycolytic metabolism in in lung fibroblasts (14, 20, 50). While it is unknown exactly how HIF-1α is induced downstream of TGF-β, increased mitochondrial activity, including production of succinate and reactive oxygen species, has been linked with stabilization of HIF-1α protein in this context (20, 50). We found that HIF-1α-dependent transcripts are enriched in the genes regulated by mTOR inhibition. While HIF-1α may be activated downstream of mTOR-dependent increases in mitochondrial activity, mTORC1 has also been shown to increase HIF-1α protein through increased cap-dependent translation of HIF-1α protein (24, 27, 47, 51). Further experiments will be required to determine the relative contribution of these mechanisms to TGF-β-induced HIF-1α activation and glycolytic reprogramming in lung fibroblasts.

Our analysis of fibroblast gene expression from patients with pulmonary fibrosis suggests that similar upregulation of amino acid biosynthesis, glycolysis, and oxidative phosphorylation occur *in vivo*. We found that within each fibroblast population, expression of mTOR and ATF4-dependent transcripts was increased in cells from pulmonary fibrosis patients. Across populations, expression of mTOR and ATF4-dependent transcripts was highest in disease-specific PLIN2^+^ and HAS1^High^ populations. Fibroblast populations, which contain the most cells from control donors, exhibited the lowest expression. While our understanding of the locations and roles of the various fibroblast populations increases, our current work provides evidence that metabolic reprogramming may contribute to the phenotypes of each fibroblast subpopulation.

## Author contributions

Conception and design: RBH, GMM

Acquisition of data: RBH, KWDS, EO, JCHS, AYM, YT, ORS PSW

Analysis and interpretation of data: RBH, KWDS, MVA, RCA, MEG, GMM

Manuscript writing: RBH, KWDS, GMM

Final approval of manuscript: KWDS, MVA, RCA, EMO, MEG, JCHS, PSW, AYM, ORS, YT, GMM, RBH

## Funding

R01ES010524 and W81XWH-22-1-0787 (GMM), and R01HL151680 (RBH).

## Notes

### Competing Interest Statement

The authors have declared no competing interest.

