## Supplemental Data File for "ATF4 and mTOR regulate metabolic reprogramming in TGF-β-treated lung fibroblasts"

Figure S1

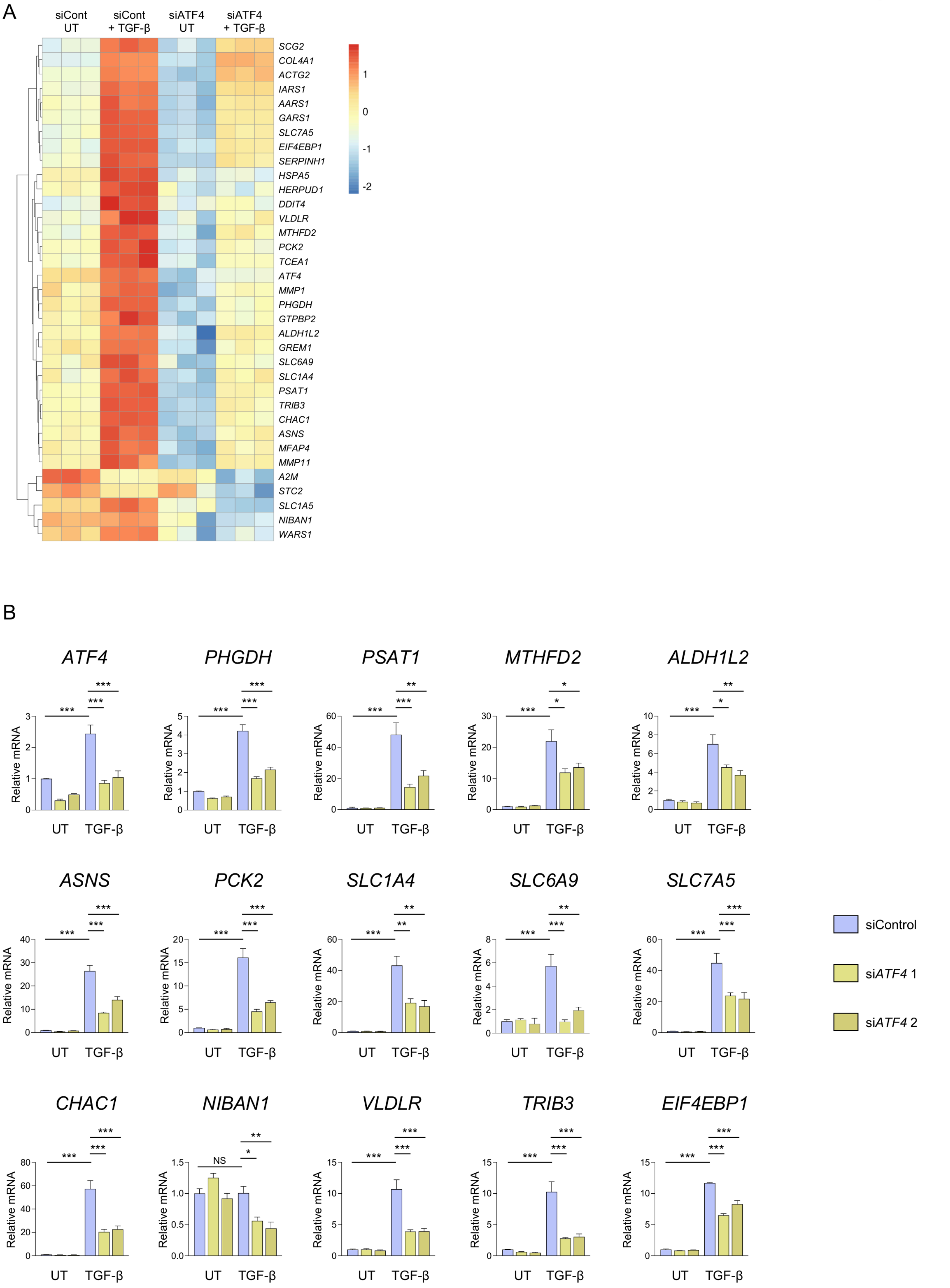
A
siCont UT
siCont + TGF-β
siATF4 UT
siATF4 + TGF-β
SCG2
COL4A1
ACTG2
IARS1
AARS1
GARS1
SLC7A5
EIF4EBP1
SERPINH1
HSPA5
HERPUD1
DDIT4
VLDLR
MTHFD2
PCK2
TCEA1
ATF4
MMP1
PHGDH
GTPBP2
ALDH1L2
GREM1
SLC6A9
SLC1A4
PSAT1
TRIB3
CHAC1
ASNS
MFAP4
MMP11
A2M
STC2
SLC1A5
NIBAN1
WARS1
1
0
-1
-2
B
ATF4
PHGDH
PSAT1
MTHFD2
ALDH1L2
ASNS
PCK2
SLC1A4
SLC6A9
SLC7A5
CHAC1
NIBAN1
VLDLR
TRIB3
EIF4EBP1
Relative mRNA
UT
TGF-β
NS
siControl
siATF4 1
siATF4 2

**Figure S1. ATF4 regulates the expression of amino acid transporters and biosynthetic enzymes in HLFs.** **(A)** Heatmap analysis of all significant DEGs ($-Log_{10}FDR \leq 0.05$) between TGF-β-treated HLFs with either control or ATF4 knockdown. **(B)** qRT-PCR analysis of expression of ATF4 target genes in control and ATF4 knockdown cells cultured in the presence or absence of TGF-β. *$P<0.05$, **$P<0.01$, ***$P<0.001$.

Figure S2

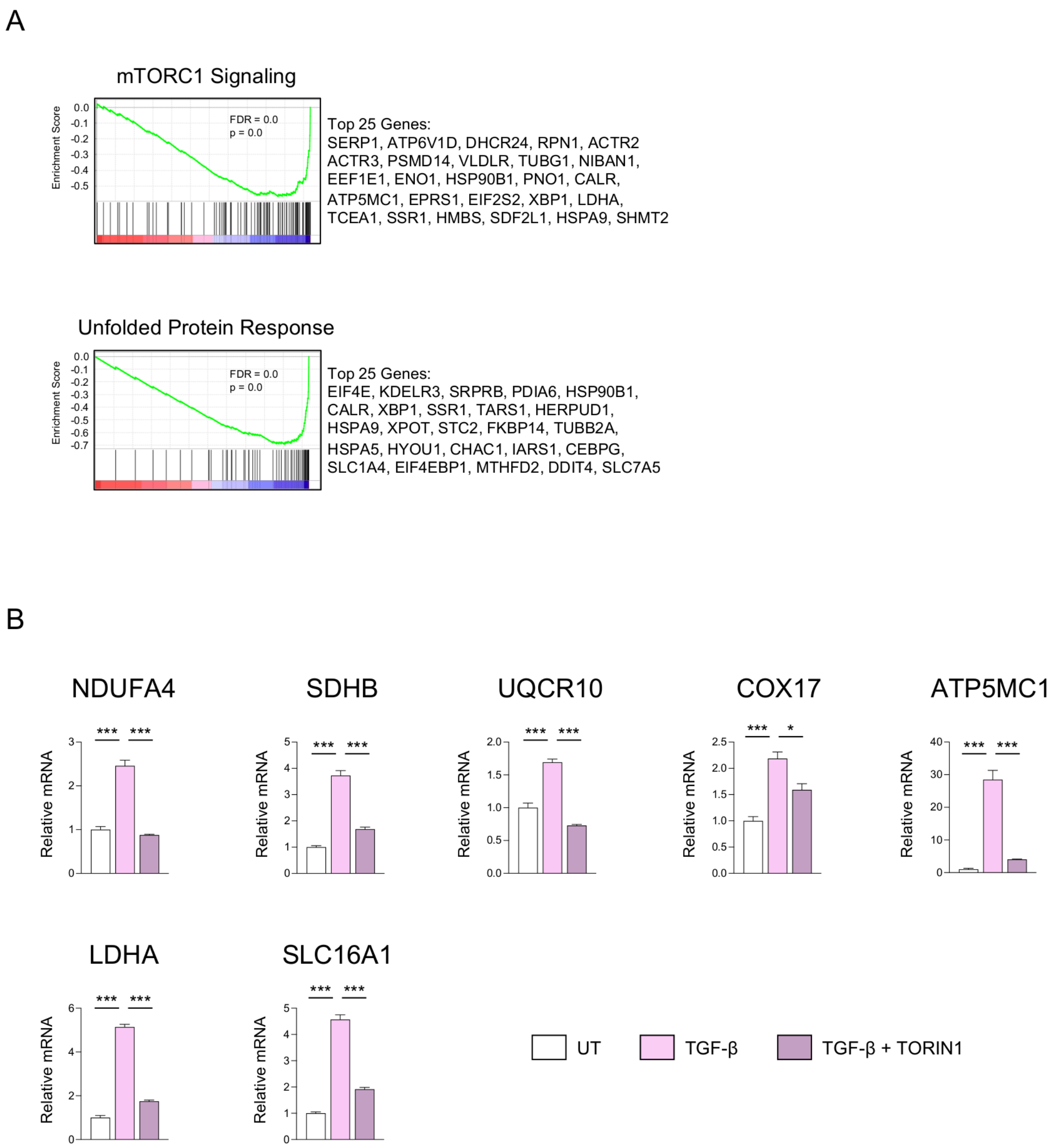


**Figure S2. mTOR regulates the expression of genes encoding glycolytic enzymes and subunits of the mitochondrial respiratory chain. (A)** Enrichment plots of significantly enriched metabolism-related Hallmark pathways regulated by mTOR inhibition in TGF-β-treated HLFs. The top 25 enriched genes in each pathway are indicated. **(B)** qRT-PCR analysis of expression of glycolytic and mitochondrial respiratory chain genes in HLFs treated with TGF-β in the presence or absence of TORIN1. **P*<0.05, ***P*<0.01, ****P*<0.001.

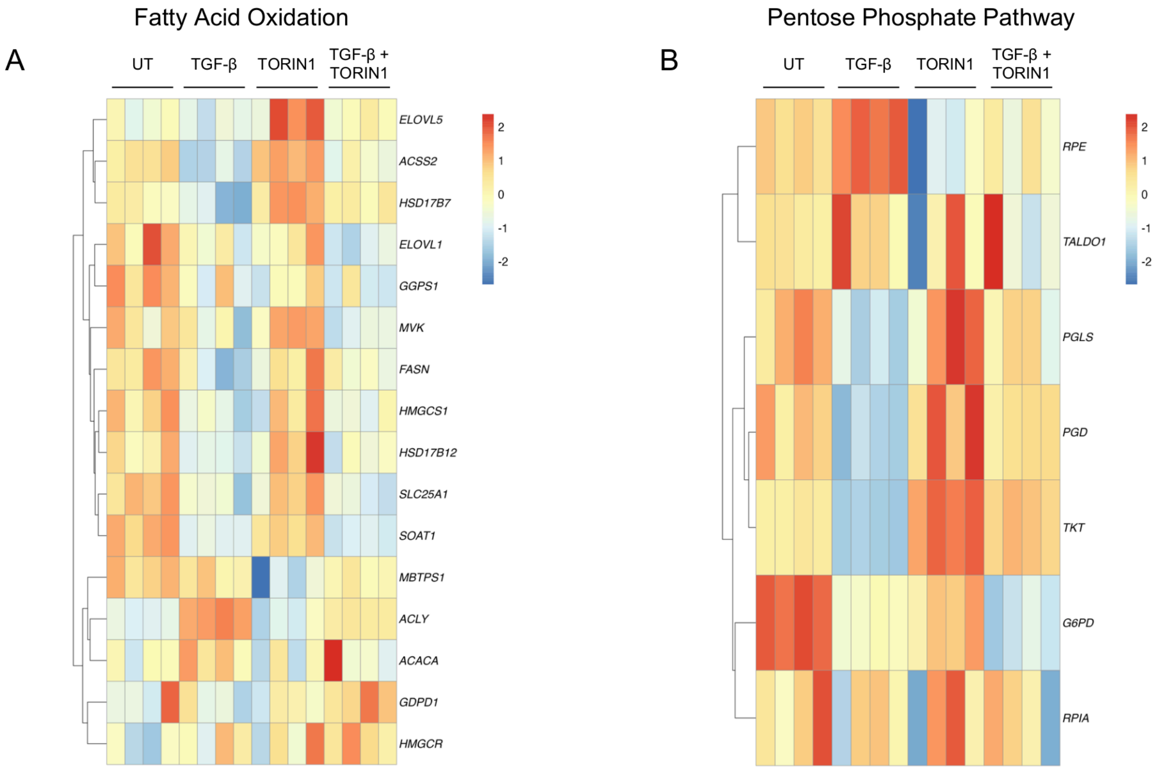


**Figure S3. mTOR inhibition does not significantly affect gene expression of fatty acid metabolic enzymes or pentose phosphate pathway enzymes (A)** Heatmap analysis of expression of genes involved in fatty acid metabolism in HLFs treated with TGF-β in the presence or absence of TORIN1. **(B)** Heatmap analysis of expression of genes encoding pentose phosphate pathway enzymes in HLFs treated with TGF-β in the presence or absence of TORIN1.

Figure S4

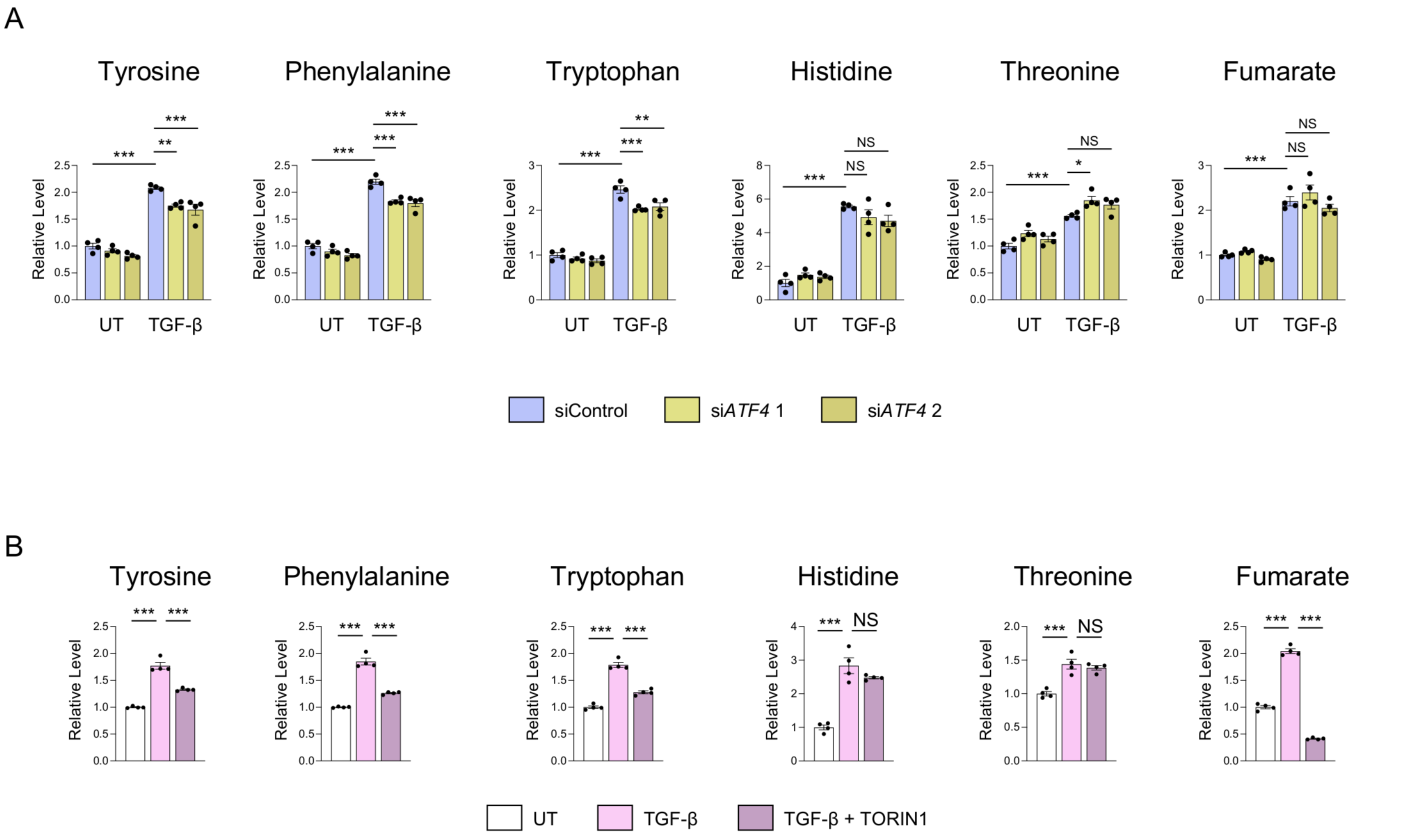


**Figure S4. ATF4 and mTOR regulate TGF-β-induced changes in cellular amino acid levels in HLFs. (A)** Relative cellular levels of the indicated amino acids in HLFs treated with TGF-β for 48 hours. Cells were transfected with either nontargeting siRNA or siRNA targeting ATF4 **(B)** Relative cellular levels of the indicated amino acids in HLFs treated with TGF-β for 48 hours. Cells were cotreated with TORIN1 as indicated. *$P$<0.05, **$P$<0.01, ***$P$<0.001.

Figure S5

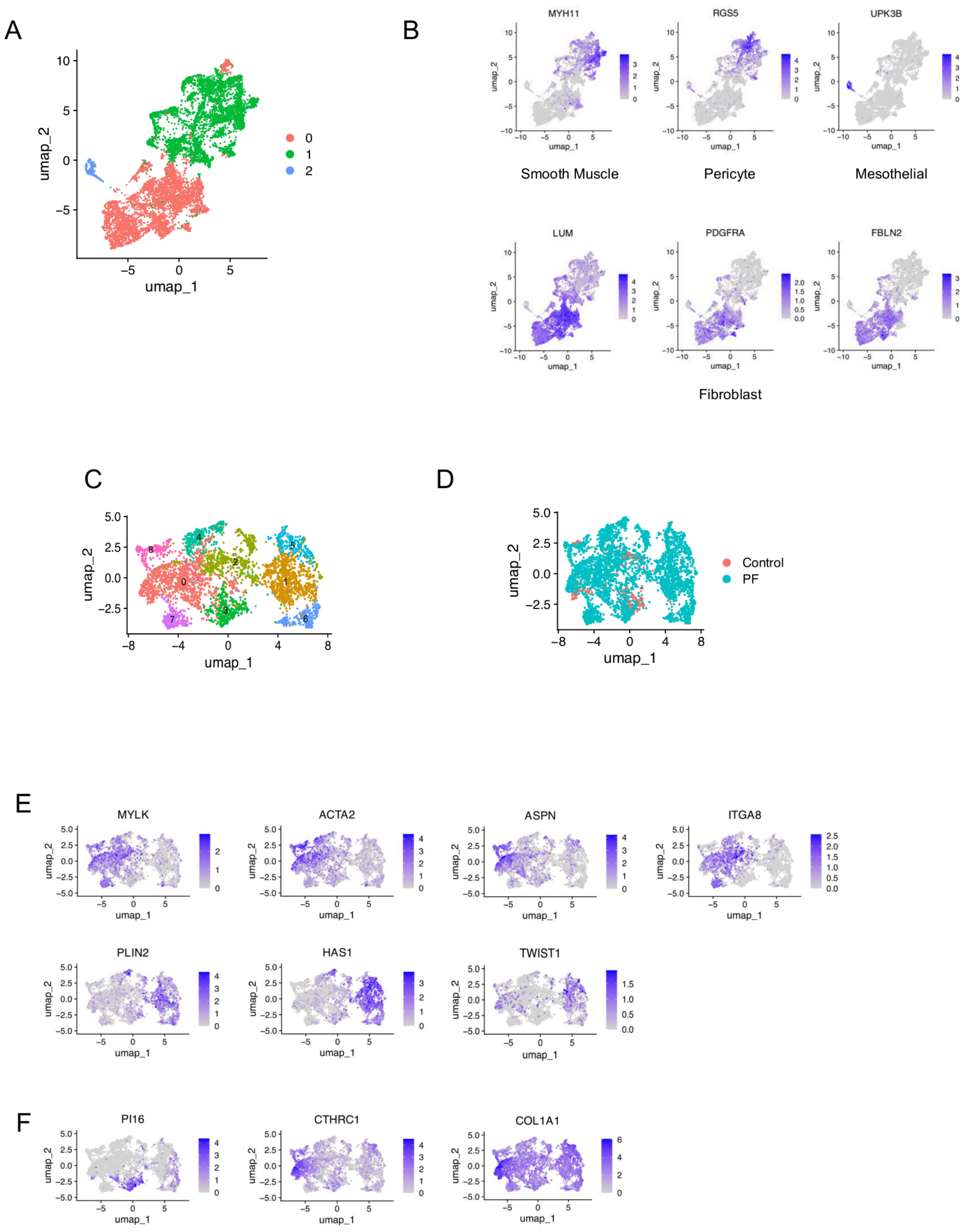
A
umap_2
umap_1
0
1
2
B
MYH11
RGS5
UPK3B
Smooth Muscle
Pericyte
Mesothelial
LUM
PDGFRA
FBLN2
Fibroblast
C
D
Control
PF
E
MYLK
ACTA2
ASPN
ITGA8
PLIN2
HAS1
TWIST1
F
PI16
CTHRC1
COL1A1

**Figure S5. Single cell characterization of mesenchymal cells from control and pulmonary fibrosis lungs.** **(A)** Clustering of *COL1A1*, *PDGFRA*, *PDGFRB*, and *LUM* expressing mesenchymal cells from 10 control lungs and 20 lungs from pulmonary fibrosis patients. **(B)** UMAP projections of expression of smooth muscle, pericyte, mesothelial cell, and fibroblast markers in the mesenchymal populations from (A). **(C)** Subclustering of fibroblast populations from cluster 0 from (A). **(D)** UMAP stratification of fibroblasts from control and pulmonary fibrosis lungs **(E)** UMAP projection of expression of genes used to assign fibroblast subpopulations as defined by Habermann et al {Habermann, 2020 #1006}. Myofibroblasts (*MYLK*, *ACTA2*), PLIN2+ Fibroblasts (*PLIN2*), HAS1High Fibroblasts (HAS1$^{High}$, PLIN2$^{Low}$, TWIAT1$^{+}$). Expression of additional myofibroblast markers ITGA8 and ASPN is also shown. (G) respiratory chain subunits in fibroblasts from pulmonary fibrosis lungs.

**Supplementary Materials and Methods**

**Primers used for qRT-PCR:**

*ATF4*- F: 5’-TTAAGCCATGGCGCTTCTCA-3’, R: 5’-GCTGGAATCGAGGAATGTGC-3’

*PHGDH*- F: 5’-GGAGGAGATCTGGCCTCTCT-3’, R: 5’-GTCATTCAGCAAGCCTGTCG-3’

*PSAT1*- F: 5’-GCGGCCATGGAGAAGCTTAG-3’, R: 5’-ATGCCTCCCACAGACACGTA-3’

*MTHFD2*- F: 5’-AAACACATCTGTCTGGTATGGT-3’, R: 5’-TGGTTAGGTCACAACTAGGAGTC-3’

*ALDH1L2*- F: 5’-GCACTAATTGGCCAGAGCCT-3’, R: 5’-AGCCAGAGGGTCAGCTTTTC-3’

*ASNS*- F: 5’-CCCCGATTTACTGGCTGCTA-3’, R: 5’-GAGCCTGAATGCCTTCCTCA-3’

*PCK2*- F: 5’-GTGCCATGGCCGCATTG-3’, R: 5’-TCTGGATGCTACGGCATGAT-3’

*SLC1A4*- F: 5’-CCCAAAGAGACGGTGGACTC-3’, R: 5’-TACGGAAAGCTGCAACCACA-3’

*SLC6A9*- F: 5’-CTTCCCCAGAACAGAATGGTG-3’, R: 5’-CGATCTGGTTGCCCCAGTTG-3’

*SLC7A5*- F: 5’-TGATCATCCTGCTGGGCTTC-3’, R: 5’-CCACATCCAGTTTGGTGCCT-3’

*NIBAN1*- F: 5’-TCTCTCGCCTCTCGAAGGAA-3’, R: 5’-CTCCTGGAGGTTGATCCGAC-3’

*VLDLR*- F: 5’-CCAGTGTCCTGGAGATGTGA-3’, R: 5’-GGTGAACTCGTCGGGACTAC-3’

*TRIB3*- F: 5’-GCCCACGCGGAACGA-3’, R: 5’-TTCTTCCTGGACAGGGAACC-3’

*EIF4EBP1*- F: 5’-GATCATACTGGGCAGGCGTT-3’, R: 5’-GCTGGTGTTCACGAAGAGGA-3’

*NDUFA4*- F: 5’-TCTCTTGCGTCTGGCATTGT-3’, R: 5’-TGGGACCCAGTTTGTTCCAG-3’

*SDHB*- F: 5’- CACTCTAGCTTGCACCCGAA-3’, R: 5’-ACATGTGTGGAAGAGGGTAGA-3’

*UQCR10*- F: 5’-ATCGTGGGCGTCATGTTCTT-3’, R: 5’-CAGCTTCCCCTCGTTGATGT-3’

*COX17*- F: 5’-GTGGTCGGGTCTCTGTTGAC-3’, R: 5’-AAGCTTGCCGTTCTCCTCTC-3’

*ATP5MC1*- F: 5’-GTGGTCGGGTCTCTGTTGAC-3’, R: 5’-AAGCTTGCCGTTCTCCTCTC-3’

*LDHA*- F: 5’-TCCGGATCTCATTGCCACG-3’, R: 5’-AGCTGATCCTTTAGAGTTGCCA-3’

*SLC16A3*- F: 5’-TCCGGATCTCATTGCCACG-3’, R: 5’-AGCTGATCCTTTAGAGTTGCCA-3’

*RPL13*- F: 5’-GTCGTACGCTGTGAAGGCAT-3’, R: 5’-GGAAAGCCAGGTACTTCAACTT-3’
